# LoRTIS Software Suite: Transposon mutant analysis using long-read sequencing

**DOI:** 10.1101/2022.05.26.493556

**Authors:** Martin Lott, Muhammad Yasir, A. Keith Turner, Sarah Bastkowski, Andrew Page, Mark A. Webber, Ian G. Charles

## Abstract

To date transposon insertion sequencing (TIS) methodologies have used short-read nucleotide sequencing technology. However, short-read sequences are unlikely to be matched correctly within repeated genomic regions which are longer than the sequence read. This drawback may be overcome using long-read sequencing technology. We have developed a suite of new analysis tools, the “LoRTIS software suite” (LoRTIS-SS), that produce transposon insertion site mapping data for a reference genome using long-read nucleotide sequence data.

Long-read nucleotide sequence data can be applied to TIS, this enables the unique mapping of transposon insertion sites within long genomic repeated sequences. Here we present long-read TIS analysis software, LoRTIS-SS, which uses the Snakemake framework to manage the workflow. A docker image is provided, complete with dependencies and ten scripts are included for experiment specific data processing before or after use of the main workflow. The workflow uses long-read nucleotide sequence data such as those generated by the MinION sequencer (Oxford Nanopore Technologies). The unique mapping properties of long-read sequence data were exemplified by reference to the ribosomal RNA genes of *Escherichia coli* strain BW25113, of which there are 7 copies of ∼4.9 kbases in length that are at least 99% similar. Of reads that matched within rRNA genes, approximately half matched uniquely. The software workflow outputs data compatible with the established Bio-TraDIS analysis toolkit allowing for existing workflows to be easily upgraded to support long-read sequencing.

## Introduction

Transposon insertion sequencing (TIS) technologies may be used to assay simultaneously every gene in the bacterial genome to find genotype-phenotype associations. However, all previous TIS methods have used sequencing platforms that generate relatively short nucleotide sequence reads of hundreds of bases, and this has limitations caused by transposon insertion sites within repeated nucleotide sequences. Thus, sequence reads that map with transposable elements and ribosomal RNA operons cannot be located uniquely if the sequence read is shorter than the length of repeat, and there is a lack of sequence diversity between the repeating units. Thus, informative genomic phenomena associated with specific repeat units may be missed. Advances in nucleotide sequencing technology have led to devices which are both portable and capable of producing sequence reads of thousands of bases. We present a new bioinformatic software package which uses long-read nucleotide sequence data (Yasir et al. 2022), such as that generated by the widely available Oxford Nanopore Technologies “MinION” sequencer, for TIS. The use of longer sequence reads has the additional advantage of allowing the unique assignment of transposon insertion sites to one unit of repeated nucleotide sequences in the bacterial genome. However, the longer sequence reads have a higher error rate, potentially complicating the matching process. This was exemplified with the *E. coli* ribosomal RNA (rRNA) operons, of which there are 7 heterologous copies of about 4.9 kb. We quantified the nucleotide sequence divergence within these long repeated sequences and show that long-read nucleotide sequence data generated from a MinION sequencer unambiguously mapped transposon insertion sites to a single rRNA operon. Our software provides data handling from the FAST5 data files generated by the MinION sequencer through to insertion site plot files which can be visualised in a genome browser such as *Artemis* (Carver et al. 2012). The software package uses the Snakemake (Mölder et al. 2021) framework for workflow management and is provided as a docker image which includes all dependencies and can be deployed by a single command

## Methods

LoRTIS-SS and supporting documentation, including examples, are publicly available from the Quadram Institute Bioscience GitHub repository (https://github.com/quadram-institute-bioscience/LoRTIS/). The recommended method of installation is Docker as containerisation allows the various dependencies to be seamlessly imported on a Linux system, such as an MRC CLIMB instance (Connor et al. 2016). At its core is a novel Snakemake workflow which implements all aspects of matching long- and short-read nucleotide sequence data to a reference genome nucleotide sequence.

### Utilities for the pre-processing of nucleotide data

Two scripts are provided which allow like-for-like comparisons to existing short-read TIS experiments. Firstly, a script is provided to randomly subsample the reads and produce a new FASTQ sequence data file with a user-specified number of reads. Next, to compare the effectiveness of the long and short read approaches, the “shred-reads.py” script is provided to split long-read sequences into shorter ones.

### Matching nucleotide sequence reads to the reference genome nucleotide sequence

Existing Bio-TraDIS (Barquist et al. 2016) software uses BWA (Li 2013) or SMALT (Ponstingl and Ning 2015) to match short-read nucleotide sequences to those of a reference genome. If a match is found at more than one location, as occurs within repeated genomic sequences, then there is a Bio-TraDIS option to choose one of the matching locations randomly, or to discard the match. For long-read sequences, the LoRTIS-SS workflow takes FAST5 format data files from the MinION sequencer and generates FASTQ files using the Guppy Basecalling software (version 3.6.0) running in High Accuracy Calling (HAC) mode. QCat (version 1.1.0) is imported using Conda and run twice, firstly, using the standard configuration, to demultiplex the FASTQ reads according to barcodes, then a second time with transposon sequence information to trim transposon sequences from the reads. The trimmed reads are matched to a reference genome nucleotide sequence using Minimap2 (Li 2018) (version 2.17-r941) to generate plot files in an analogous manner to short-reads data. The error rate for Nanopore sequencing has been estimated at 1%-3% (Delahaye and Nicolas 2021) which is considerably higher than that for Illumina, making the BWA or SMALT alignment methods unsuitable. Minimap2 finds matches between sequences with differences of up to 15% and is therefore an appropriate matching tool to use when higher levels of sequencing errors are expected. In contrast to Bio-TraDIS, when there is more than one mapping location, the sequence read is discarded. LoRTIS-SS then uses the Minimap2 output to generate transposon insertion site plots, which can be viewed alongside an annotated genome in a genome browser such as *Artemis (Carver et al. 2012)*, and processed further using Bio-TraDIS to provide statistical data for each annotated gene. Thus, by taking this approach, the greater error rate of nanopore generated sequence reads compared to short-reads sequences are accounted for intrinsically.

### Post-Processing

A further eight scripts are provided to aid understanding of the insertion plots. Firstly, script “change-plot.py” is provided to change the sign of insertions on the reverse strand such that the number of insertions will be displayed below the axis in *Artemis*. Secondly, the “combine-plot-files.py” script is provided to combine plot files, as may be required, providing backwards compatibility with Bio-TraDIS. Furthermore, new software functions include the flagging of very short genomic regions with more than 5 insertions at a position (“spikes.py”). These sites, often within coding sequences, can represent transposon insertion mutants with significant selective advantage and are therefore regions of interest that existing versions of Bio-TraDIS did not flag. In addition, a script, “operons.py,” takes as input a user-provided reference genome annotation in EMBL format and contextualizes genes in genome order to provide putative operon information to assist with TIS data interpretation. TIS data will often demonstrate a low level of reads across the genome, including within genes that are known to be essential. These matched reads may originate from non-viable mutants that remain present in the transposon mutant library as whole cells or as DNA from dead and lysed cells. This “background” signal may interfere with the calling of candidate essential genes, especially when relatively large numbers of reads are matched with a reference genome. The “remove-background-insertions.py” script removes this background by setting to zero any transposon insertion sites for which the matched sequence reads is below a chosen threshold, in our experience a threshold which removes fewer than 3 insertions per site will suffice. The “list-non-empty-sites.py” script provides a list of genome positions where there are insertions sites, as a csv file. Finally, the “reduce-insertion-plot.py” works in similar fashion to shuffle-reads.py but instead of subsampling the reads, this randomly subsamples insertions in the insertion plot so they can be compared like-for-like across experiments. Results can then be supplied to AlbaTraDIS (Page et al. 2020) for large scale comparative analysis between experimental conditions.

## Results

### Nucleotide sequence divergence between the 7 copies of rRNA gene clusters

As a robust test, the rRNA (rrs/rrl) gene operons were chosen to determine the feasibility of matching long-read sequences uniquely to repeated genomic nucleotide sequences. At approximately 4.9 kb, these operons are the longest repeated genetic elements in the *E*.*coli* BW25113 genome. There are 7 copies (A, B, C, D, E, G, H), and they all include the genes coding for the 16S and the 23S rRNA which are about 1.5 kb and 2.9 kb in length respectively (Fig 1). These genes are at least 99% identical between different operons and are separated by a more divergent sequence of approximately 0.4kb which incorporates one or two sometimes different tRNA genes, depending on the operon. Alignment of the operons using Clustal Omega (Sievers et al. 2011) identified the heterogeneous base positions, and demonstrated that there is a region of approximately 500 bp which are identical in all 7 copies (Fig1). Therefore, to identify transposon insertion sites uniquely, the sequence reads need to be substantially longer than this.

**Fig. 1.**
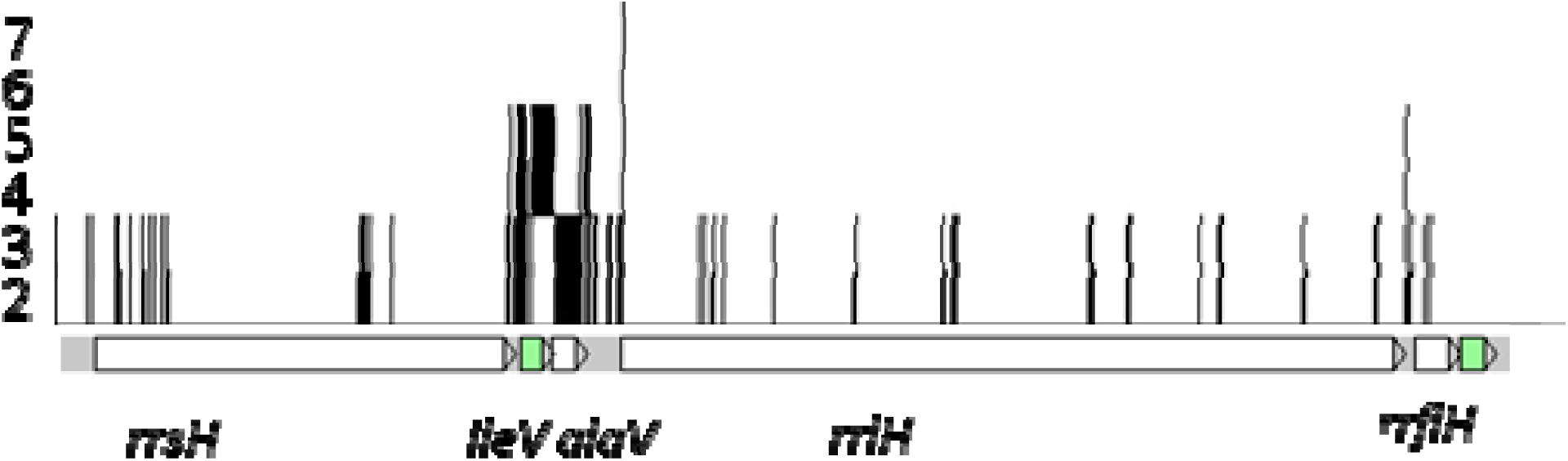
Polymorphic regions within the ribosomal RNA genes. A genetic map of the rrsH-rrlH rRNA operon incorporating the *ileV* and *alaV* tRNA genes is shown alone the bottom of the figure. Vertical bars above this indicate the locations of differences between the operons and the *y-*axis indicates the number of operons at that site which have a different base from that which is most common. The *ileV* and *alaV* differ between the operons, but most of the differences are single or a small number of bases. In the *rrsH* gene there are approximately 500 contiguous bases which are identical between all 7 rRNA operon copies

### Identifying transposon insertion sites uniquely within genomic repeating nucleotide sequences using long-read sequences

To identify unique matches within repeated genomic nucleotide sequences a sequence read must include sufficient unique heterogeneous sequences either adjacent to or within the repeats. Therefore, short sequence reads used previously for TIS which are tens of bases in length will not match uniquely within much of the rRNA operons. This can be circumvented by using long-read sequence data, and our long-read datasets included sequence reads of up to 10 kb and the majority were over 500 bp in length. Thus, a significant number of these long-reads are likely to match across the heterogeneous sequences of repeats allowing for a unique match. However, nanopore-generated sequence reads have an error rate of up to 3%, which is far greater than the short-read sequences that have been used traditionally for TIS. So, even unique matches are only likely to have a nucleotide sequence similarity of approximately 97% using long-read nanopore-generated sequences. Comparing different criteria for when a read should be kept, we found that excluding reads which do not map uniquely allows us to retain over 97% of all reads which map to the reference genome.

### Comparison with existing software

In general, bacteria have a number of regions of repeated nucleotide sequences within their genomes, which makes it impossible to uniquely map transposon insertion sites using sequence reads that are shorter than the unit of sequence repeat unless the repeating units show heterogeneity. Where there is ambiguity, existing software typically will pick one of the matching positions at random. Using long nucleotide sequence reads allows transposon insertion sequences to be matched uniquely including within long, repeated genomic sequences. This is because long-reads span across greater distances, increasing the likelihood of spanning heterogeneous sequences, or by including the unique sequences that flank the repeats. Even so, some long-reads do not match uniquely, and these are discarded. But the long-read sequence data includes sufficient reads to allow unique mapping. Thus, for our data we observed that approximately 50% of the reads that matched to the rRNA genes, of which there are seven copies of 4.9 kb in the *E. coli* reference genome, matched uniquely (Fig. 2). This proportion can likely be increased by improvements to the LoRTIS protocol aimed at increasing the yield of longer reads. All sequence data used for this analysis has been deposited in the European Nucleotide Archive (accession number E-MTAB-11351) and was described in detail in (Yasir et al. 2022).

**Fig. 2.**
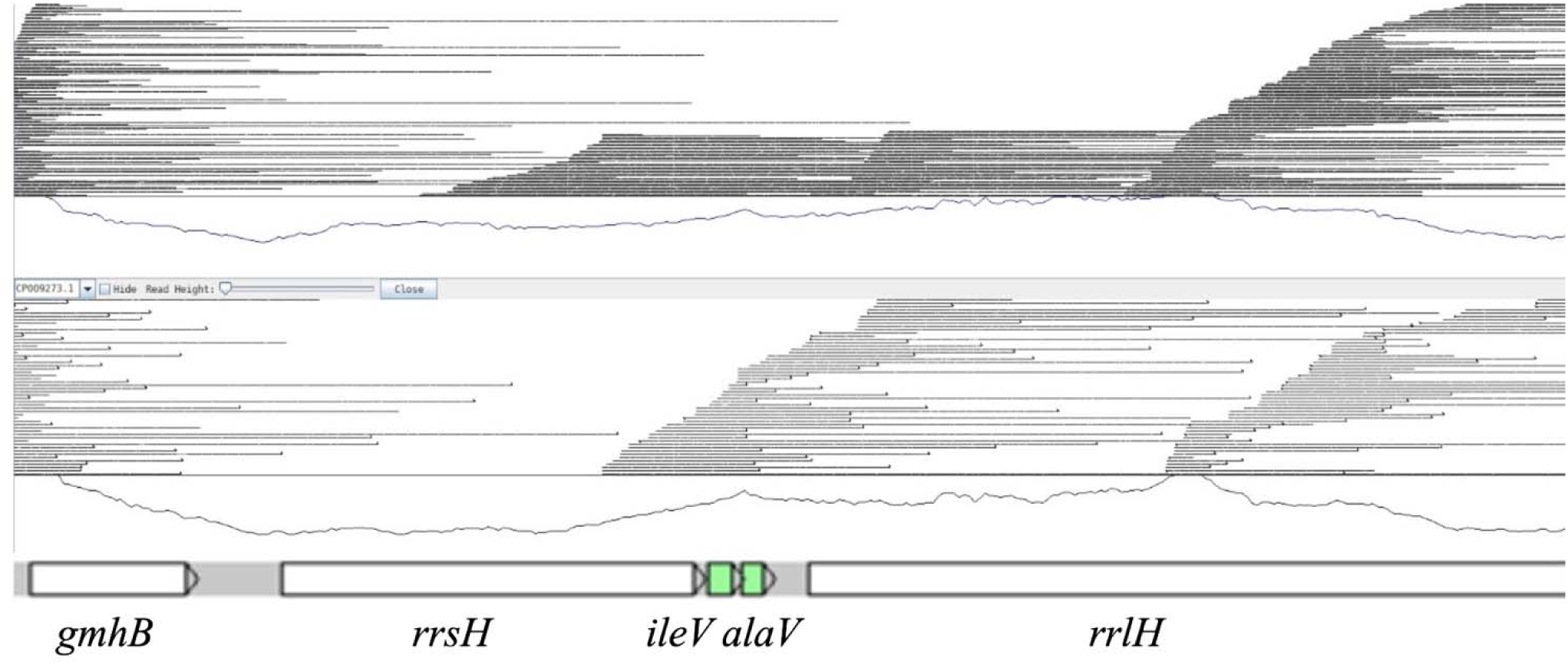
Locations of long sequence reads that matched uniquely to the *rrsH-rrlH* rRNA gene operon of *E. coli*. A genetic map of the *rrsH-rrlH* rRNA gene operon is presented, above which are depicted the locations and lengths of uniquely matched long nucleotide sequence reads (fine horizontal lines). The upper and lower panels depict long-read sequences from two different replicate experiments, and the lower trace in each is a measure of the relative coverage.

## Conclusion

The LoRTIS technology and custom software suite may thus provide insights around repeated genomic sequences that other TIS technologies miss.

